# Organ-specific transcriptomic and proteomic changes in thermomorphogenesis in *Arabidopsis thaliana* seedlings

**DOI:** 10.1101/2025.08.05.668523

**Authors:** Jingmin Hua, Shitij Gupta, Dominique Jacques-Vuarambon, Rodrigo S. Reis

**Affiliations:** Institute of Plant Sciences, University of Bern, Bern, Switzerland

**Keywords:** thermomorphogenesis, warm temperature, gene expression, whole seedling, organ-specific

## Abstract

Plant acclimatisation to mild, yet prolonged temperature increases is critical for survival amid global warming. Temperature increases within the physiological range—24-30°C, for the model *Arabidopsis thaliana* Col-0—promote coordinated response across the different plant organs, termed thermomorphogenesis. However, most transcriptomic studies analyse whole-seedling material, hindering insights into organ-specific responses. Here, we analysed public whole-seedling transcriptomes and produced transcriptomic and proteomic data for root, hypocotyl, and cotyledon in Arabidopsis under warm temperature (28°C vs 21°C). Although independent whole-seedling datasets showed broad rank-order concordance, standard thresholding for differentially expressed genes (DEGs) yielded limited cross-study overlap. We compared DEGs in whole seedling and dissected organs and observed that 67% of up-regulated and 78% of down-regulated genes in whole seedling are not DEGs in any individual organ, evidencing a need for organ-specific data. We then found that warm temperature mostly suppressed protein accumulation in hypocotyl, likely through translational inhibition. Furthermore, comparison of differentially expressed proteins (DEPs) and DEGs revealed widespread post-transcriptional regulation and transcriptional buffering. We identified several promising genes still uncharacterized for a function in thermomorphogenesis, such as *DRG1-3* (up DEG in all organs and seedling), *MSRB6* (up DEG in all organs and seedling, but down DEP in root and hypocotyl), *AGL20*/*SOC1* (down DEG in all organs and most seedling datasets), and *TPX2*/*PRXIIC* (up DEP and DEG in root and most seedling datasets). Our organ-specific resource presents a detailed view on plant response to warm temperature and open new questions and research avenues in thermomorphogenesis.

**SIGNIFICANCE STATEMENT:** To cope with warmer conditions, plants use distinct mechanisms above and below ground to adjust the growth of their roots, stems, and leaves. Our transcriptomic and proteomic resource for Arabidopsis thaliana root, hypocotyl, and cotyledon reveals how each organ regulates mRNA and protein levels, opening new questions in plant response to warm temperatures.

## INTRODUCTION

Plants acclimatise to mild increases in ambient temperature through a coordinated process termed thermomorphogenesis—a suite of non-stressful morphological and physiological changes (Casal and Balasubramanian, 2019). This thermal response is regulated by a complex network of molecular thermosensors and signalling pathways (Yadav *et al*., 2025). Perception of warming temperatures involves multiple distinct mechanisms, including phytochrome B (phyB) thermal reversion from its active Pfr form to its inactive Pr state (Legris *et al*., 2016; Jung *et al*., 2016), EARLY FLOWERING 3 (ELF3) temperature-dependent liquid-liquid phase separation into inactive biomolecular condensates (Jung *et al*., 2020), and an RNA thermoswitch in the 5’-untranslated region (UTR) of *PHYTOCHROME-INTERACTING FACTOR 7* (*PIF7*) mRNA that relaxes at warmer temperatures to enhance translation (Chung *et al*., 2020). Most warm temperature sensing mechanisms converge on PHYTOCHROME-INTERACTING FACTOR 4 (PIF4), a central bHLH transcription factor in plant growth (Kerbler and Wigge, 2023), in which warm temperatures stabilize PIF4, allowing for its induction of auxin biosynthesis genes (Franklin *et al*., 2011). Auxin signalling is then integrated with other phytohormones through mechanisms such as the brassinosteroid (BR) transcription factor BRASSINAZOLE-RESISTANT 1 (BZR1) binding to *PIF4* promoter and physically interacting with PIF4 in a self-reinforcing positive feedback loop (BZR1–PIF4–auxin–BR) modulated by co-factors like HIGH NITROGEN INSENSITIVE 9 (HNI9/IWS1) and BEL1-LIKE HOMEODOMAIN 1 (BLH1) (Chen *et al*., 2026; Qin *et al*., 2026; Ibañez *et al*., 2018), while the gibberellin (GA)-repressing DELLA proteins are degraded under warm conditions to relieve growth inhibition (Camut *et al*., 2019).

Thermomorphogenesis is organ-specific and tightly regulated across different plant tissues. For instance, in the embryonic stem (hypocotyl), warm temperatures trigger organ elongation driven by auxin-mediated cell expansion (Gray *et al*., 1998). In cotyledons, warmth restricts blade expansion and lowers stomatal density through PIF4-dependent suppression of cell cycle and stomatal lineage regulators such as SPEECHLESS (SPCH) (Saini *et al*., 2022; Lau *et al*., 2018). Further, cotyledons act as the primary sites of temperature perception and auxin synthesis—necessary for driving the hypocotyl elongation (Bellstaedt *et al*., 2019). Leaves, on the other hand, exhibit hyponasty (upward movement) associated with petiole elongation, creating an open rosette structure (Casal and Balasubramanian, 2019). Hyponastic movement is driven by PIN-FORMED (PIN3)-and PINOID-dependent directional auxin transport toward the abaxial side of the petiole, whereas leaf blade area is simultaneously restricted through PIF4–TCP4 transcription factor interactions that induce the cell cycle inhibitor KIP-RELATED PROTEIN 1 (KRP1) to suppress cell proliferation, thereby elevating only the abaxial side of the petiole (Saini *et al*., 2022; Park *et al*., 2019). Below ground, primary roots elongate in response to moderate warmth presumably via autonomous root-sensing pathways that function independently of shoot thermosensors (Ai *et al*., 2023; Borniego *et al*., 2022). Unlike elongation driven by cell expansion in the hypocotyl and petiole, root thermomorphogenesis is driven mainly by increased cell division rates in the root apical meristem via cell-autonomous auxin biosynthesis and PIN-mediated auxin gradients (Ai *et al*., 2023), together with cortical cell expansion that optimizes root carbon partitioning between biomass formation and respiration (Colombi *et al*., 2026).

In an ecological and agronomical context, thermomorphogenesis is an acclimatisation process that leverages phenotypic plasticity to cope with warmer conditions, while leading to negative consequences to crop yield. For instance, the open rosette architecture, hyponasty, and petiole elongation increase ventilation and cooling via transpiration, lowering temperatures and risks of heat damage to photosynthetically active structures (Lee *et al*., 2020). This coordinated response also includes primary root elongation to enable plants to reach deeper, cooler, and moister soil layers, given that warming is often associated with reduced water availability in the soil (Hayes *et al*., 2025). Warm temperatures further trigger early flowering likely to ensure that reproductive development is completed prior to possible worsening in the environmental conditions (Ludwig *et al*., 2021). However, these thermomorphogenic strategies have significant agricultural trade-offs and yield challenges for major crop species such as wheat, rice, maize, and soybean, amid global warming (Asseng *et al*., 2015; Zhao *et al*., 2017). Exacerbated stem elongation redistributes carbon resources away from storage and grain organs, producing thin, spindly plants prone to lodging from wind and rain (Lee *et al*., 2020). Furthermore, PIF4 activation leads the physiological trade-off between growth and immunity to favour growth, suppressing defence and making crops more vulnerable to pathogen attack under warmer conditions (Gangappa *et al*., 2017; Shields *et al*., 2025).

The costly phenotypical changes in response to warmer temperatures are under tight gene expression regulation—a major research focus during the past few decades. There has been growing evidence that plants employ different mechanisms to sense and respond to warm temperature in organs below (root) and above (shoot) ground (Borniego *et al*., 2022; Jacob *et al*., 2025). However, our mechanistic understanding has been largely limited to shoot, particularly, hypocotyl elongation (Delker *et al*., 2022; Quint *et al*., 2023). Furthermore, most of the studies to date that employ transcriptomic analysis have been limited to whole seedling material, hindering insights on organ-specific response to warmer temperatures. Here, we present transcriptomic and proteomic data for root, hypocotyl, and cotyledon produced with *Arabidopsis thaliana* Col-0 wild-type plants grown under commonly used experimental setup to study the response to warmer temperatures. Our comprehensive multi-omics resource will enable the community to better understand how plants coordinate their organ-specific perception and response to ambient temperature increases.

## RESULTS

### Conventional thresholding in DEG analysis yields limited cross-study overlap despite broad concordance of ranked responses

We initially compared various transcriptomic datasets produced using Arabidopsis whole seedlings treated with warm temperatures (Jin et al., 2020; Romero-Montepaone et al., 2021; Park et al., 2022; Qin et al., 2022), including our own dataset (Supporting Table 1). We produced pairwise comparisons using rank-rank hypergeometric overlap 2 (RRHO2) (Cahill *et al*., 2018) to identify overlapping signatures and visualize concordant and discordant patterns (Figure 1a). RRHO2 is a threshold-free algorithm, hence, it ranks the entire gene list by differential expression p-values and effect size direction. Overall, the datasets showed high concordance level, suggesting consistent transcriptome-wide responses to warm temperature among independent studies. However, when we compare DEGs using standard thresholds (|log2FC| ≥ 1 and adjusted p-value < 0.05), only a small DEG subset is shared by at least four out of the five datasets, regardless of whether they are up-or down-regulated (Figure 1b). It is possible that differences in treatment strength and biological variability may hinder the identification of a larger DEG overlap across studies. Indeed, experimental conditions to produce these datasets were not homogenous: temperature treatment durations were short (10 h or one day) or long (three or four days), temperature amplitude between treatment and control was in the range of 5-8°C, and light conditions varied in intensity, from 70 to 120 μmol m^−2^ s^−1^ white light (Supporting Table 1). Restricting the comparison to three datasets with the closest treatment conditions (CRA001128, CRA006077, and this work; 3-4 days at 20-21°C followed by 3-4 days at 28°C) still yielded low DEG overlap (∼4-9% for upregulated and ∼2-3% for downregulated) and high percentage of dataset-specific DEGs (∼38-51%), showing that lab-to-lab variance has a large contribution to DEG identification. Altogether, our metanalysis reveals that conventional DEG identification in thermomorphogenesis is sensitive to lab-to-lab variance and biological state heterogeneity arising from differences in treatment duration, light regime, and circadian sampling. Consequently, a third to half of reported DEGs across comparable studies were not shared—a cautionary observation for future cross-study comparisons.

**Figure 1.**
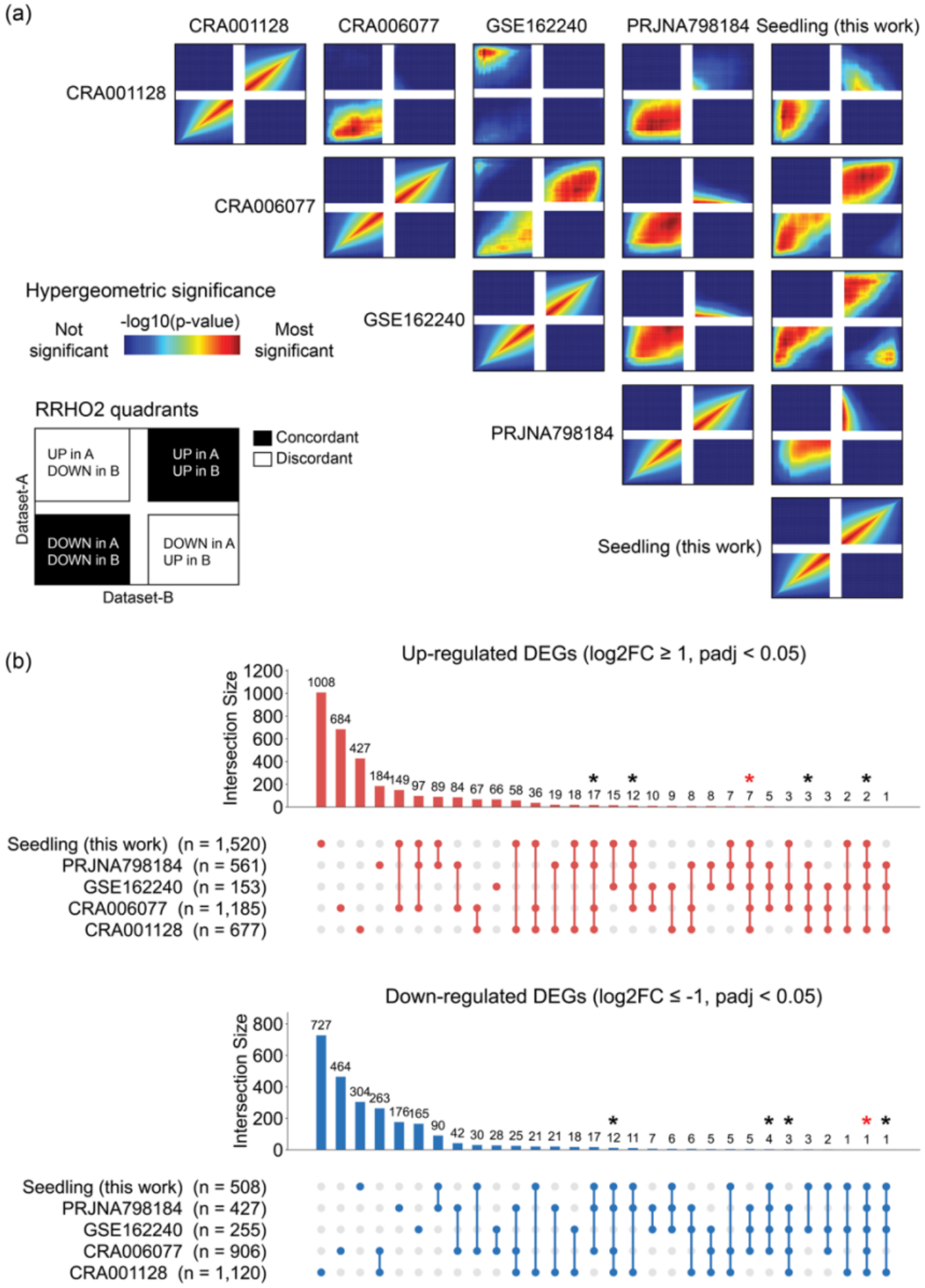
Comparison of transcriptomic responses to warm temperature across independent whole-seedling datasets in *Arabidopsis thaliana* Col-0. **(a)** Pairwise Rank-Rank Hypergeometric Overlap 2 (RRHO2) analysis comparing five seedling transcriptomic datasets. Heatmaps depict threshold-free pattern agreement across entire ranked gene lists. Colour gradient indicates hypergeometric significance (-log_10_(p-value)), ranging from non-significant (blue) to highly significant (red). Quadrants represent concordant (upper-right: UP in both; lower-left: DOWN in both) and discordant (upper-left: UP in A, DOWN in B; lower-right: DOWN in A, UP in B) gene expression directionality. **(b)** UpSet plots of differentially expressed genes (DEGs) identified using standard threshold parameters (|log_2_(fold change)| ≥ 1.0, p_adj_ < 0.05). Vertical bar charts display intersection sizes for up-regulated (top, red) and down-regulated (bottom, blue) DEGs across datasets. Total DEG counts per dataset are shown in parentheses. Asterisks highlight subsets shared across ≥ 4 datasets. Red asterisks mark core DEGs common to all five datasets.

**Table 1.** Agreement between transcriptomic and proteomic changes above statistical threshold (|log2 fold change| ≥ 1 and p-adjusted value < 0.05) in root, hypocotyl, and cotyledon after temperature treatment (28°C vs 21°C).

| Organ | AGI | Short name | Transcript (log2FC) | Transcript (padj) | Protein (log2FC) | Protein (padj) | Transcript/protein agreement |
|---|---|---|---|---|---|---|---|
| **Root** | AT5G42830 | AT5G42830 | 1.86 | 2.15E-06 | 4.13 | 2.99E-04 | concordant_G1 |
|  | AT3G21720 | ICL | 2.9 | 7.27E-14 | 4.09 | 2.36E-03 | concordant_G1 |
|  | AT1G21130 | IGMT4 | 2.6 | 1.05E-03 | 3.89 | 2.05E-04 | concordant_G1 |
|  | AT5G25450 | AT5G25450 | 1.27 | 7.34E-03 | 3.78 | 7.44E-04 | concordant_G1 |
|  | AT1G80240 | DGR1 | 2.19 | 4.42E-13 | 3.63 | 2.08E-03 | concordant_G1 |
|  | AT5G51440 | HSP23.5 | 1.63 | 1.12E-03 | 3.63 | 8.47E-03 | concordant_G1 |
|  | AT3G57330 | ACA11 | 1.09 | 1.34E-07 | 3.44 | 3.66E-03 | concordant_G1 |
|  | AT5G16010 | AT5G16010 | 1.05 | 1.53E-03 | 3.32 | 5.90E-03 | concordant_G1 |
|  | AT1G70290 | TPS8 | 1.62 | 2.64E-05 | 3.15 | 1.42E-02 | concordant_G1 |
|  | AT1G13090 | CYP71B28 | 1.19 | 2.18E-08 | 3.08 | 1.42E-02 | concordant_G1 |
|  | AT5G09290 | AT5G09290 | 1.46 | 6.53E-03 | 2.49 | 2.86E-04 | concordant_G1 |
|  | AT1G11300 | EGM1 | 1.45 | 1.09E-04 | 2.43 | 7.37E-03 | concordant_G1 |
|  | AT5G52640 | HSP90.1 | 1.94 | 3.86E-07 | 1.65 | 6.74E-03 | concordant_G1 |
|  | AT1G65970 | TPX2 | 1.96 | 9.65E-12 | 1.58 | 1.19E-03 | concordant_G1 |
|  | AT2G39310 | JAL22 | 2.01 | 1.37E-11 | 1.35 | 8.91E-03 | concordant_G1 |
|  | AT4G11650 | OSM34 | 1.89 | 2.56E-08 | 1.26 | 4.99E-02 | concordant_G1 |
|  | AT1G02920 | GSTF7 | 1.76 | 4.16E-02 | 1.25 | 1.04E-02 | concordant_G1 |
|  | AT1G02930 | GSTF6 | 2.42 | 1.72E-02 | 1.2 | 3.55E-03 | concordant_G1 |
|  | AT2G29340 | AT2G29340 | 2.05 | 1.85E-13 | 1.17 | 8.76E-03 | concordant_G1 |
|  | AT5G43330 | c-NAD-MDH2 | 1.45 | 8.88E-08 | 1.09 | 3.01E-02 | concordant_G1 |
|  | AT1G21100 | IGMT1 | 2.6 | 7.89E-19 | 1.07 | 4.80E-02 | concordant_G1 |
|  | AT2G24180 | CYP71B6 | 1.08 | 3.10E-08 | 1.07 | 3.01E-02 | concordant_G1 |
|  | AT3G01420 | DOX1 | 1.83 | 4.69E-10 | 1.03 | 3.62E-02 | concordant_G1 |
|  | AT5G52310 | LTI78 | -1.12 | 1.89E-06 | -2.15 | 1.71E-02 | concordant_G2 |
|  | AT3G16390 | NSP3 | -1.36 | 6.42E-07 | -4.72 | 1.21E-04 | concordant_G2 |
|  | AT4G02340 | AT4G02340 | -1.01 | 1.09E-06 | 3.91 | 2.86E-04 | discordant_G3 |
|  | AT1G13420 | ST4B | 1.21 | 8.63E-03 | -1.2 | 8.21E-03 | discordant_G4 |
|  | AT2G43590 | AT2G43590 | 1.17 | 1.98E-03 | -1.3 | 3.35E-02 | discordant_G4 |
|  | AT1G53680 | GSTU28 | 1.1 | 1.88E-02 | -2.58 | 2.56E-03 | discordant_G4 |
|  | AT4G04840 | MSRB6 | 1.17 | 4.27E-04 | -2.67 | 6.88E-04 | discordant_G4 |
| **Hypocotyl** | AT2G10940 | AT2G10940 | 2.83 | 3.09E-31 | 3.18 | 1.74E-03 | concordant_G1 |
|  | AT5G22580 | AT5G22580 | 2.61 | 1.98E-12 | 3.69 | 1.86E-04 | concordant_G1 |
|  | AT1G52190 | NRT1.11 | 1.74 | 2.36E-07 | 1.93 | 1.46E-03 | concordant_G1 |
|  | AT1G25230 | AT1G25230 | 1.43 | 6.88E-06 | 3.45 | 1.08E-04 | concordant_G1 |
|  | AT1G12080 | CCAP2 | 1.6 | 5.69E-04 | 1.58 | 2.38E-03 | concordant_G1 |
|  | AT4G38690 | AT4G38690 | 1.21 | 3.20E-03 | 1.09 | 1.90E-02 | concordant_G1 |
|  | AT5G25450 | AT5G25450 | 1.46 | 2.62E-02 | 2.03 | 1.10E-03 | concordant_G1 |
|  | AT2G46740 | GulLO5 | -3.52 | 1.35E-07 | -2.57 | 2.46E-04 | concordant_G2 |
|  | AT5G43350 | PHT1;1 | -1.95 | 8.84E-05 | -3.16 | 9.19E-03 | concordant_G2 |
|  | AT4G28250 | EXPB3 | -1.2 | 9.04E-04 | -2.46 | 2.85E-04 | concordant_G2 |
|  | AT1G27020 | AT1G27020 | -1.15 | 3.28E-03 | -1.45 | 1.94E-03 | concordant_G2 |
|  | AT4G04840 | MSRB6 | 1.98 | 1.84E-05 | -2.54 | 2.97E-04 | discordant_G4 |
|  | AT1G21670 | AT1G21670 | 1.07 | 1.03E-02 | -1.77 | 2.67E-02 | discordant_G4 |
| **Cotyledon** | AT2G20560 | DNAJ | 1.34 | 7.57E-03 | 5.18 | 2.02E-04 | concordant_G1 |
|  | AT5G48570 | FKBP65 | 2.3 | 1.98E-10 | 5.13 | 4.18E-03 | concordant_G1 |
|  | AT2G43620 | CHI62 | 1.65 | 5.24E-03 | 4.33 | 2.54E-04 | concordant_G1 |
|  | AT1G52000 | AT1G52000 | 2.47 | 1.65E-20 | 3.9 | 2.31E-04 | concordant_G1 |
|  | AT1G22370 | UGT85A5 | 1.19 | 1.12E-02 | 3.78 | 1.69E-03 | concordant_G1 |
|  | AT4G27860 | MEB1 | 1.14 | 2.45E-06 | 3.58 | 1.03E-02 | concordant_G1 |
|  | AT3G12580 | HSP70 | 2.2 | 5.12E-08 | 2.44 | 1.78E-04 | concordant_G1 |
|  | AT1G64400 | LACS3 | 1.05 | 4.91E-04 | 1.38 | 7.15E-03 | concordant_G1 |
|  | AT5G52640 | HSP90.1 | 2.62 | 3.60E-15 | 1.08 | 5.26E-04 | concordant_G1 |
|  | AT4G23680 | AT4G23680 | 4.1 | 5.50E-103 | 1.06 | 1.58E-04 | concordant_G1 |
|  | AT3G23830 | GRP4 | -1.02 | 5.35E-03 | -1.56 | 2.02E-04 | concordant_G2 |
|  | AT4G21770 | RLUA3 | -1.14 | 4.98E-07 | -3.53 | 3.43E-02 | concordant_G2 |

We reasoned that DEGs shared by at least four out of the five datasets are under tighter control specifically by warm temperature (Figure 2). For instance, up-regulated genes shared by all five datasets can be readily associated with heat response, e.g., heat-shock proteins *HSP70T-2* and *DNAJ* and the thermotolerance-associated transcriptional coactivator MULTIPROTEIN BRIDGING FACTOR 1C (*MBF1C*) (Suzuki *et al*., 2008), while the only such down-regulated gene is associated with cold response, i.e., *KIN1* (Chinnusamy *et al*., 2007). Most DEGs, however, are still uncharacterized for a possible role in plant response to warm temperature, such as the putative GTP-binding protein *DEVELOPMENTALLY REGULATED G-PROTEIN 1-3* (*DRG1-3*).

**Figure 2.**
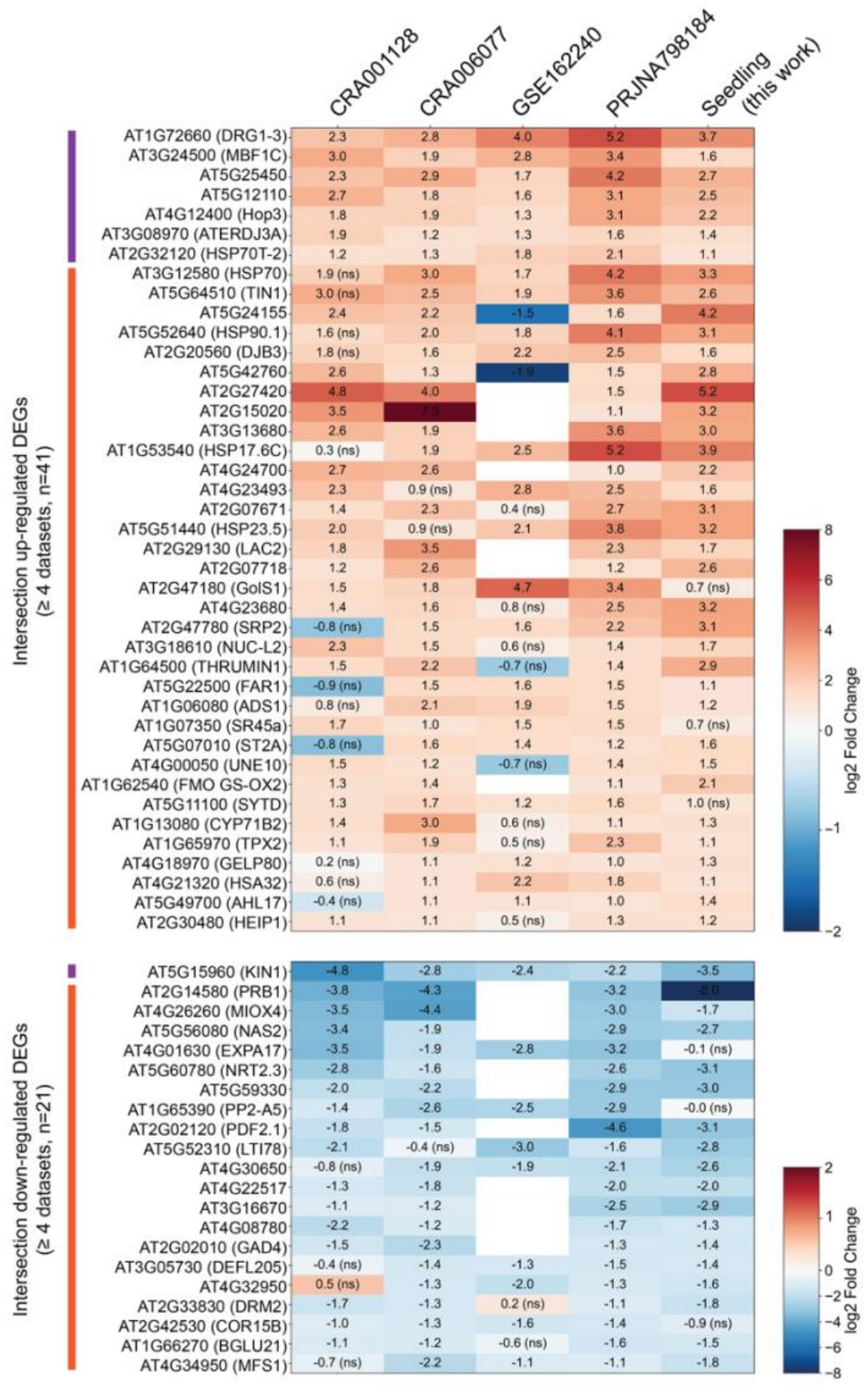
Comparison of core transcriptomic changes to warm temperature shared by independent whole-seedling datasets. Heatmap for core differentially expressed genes (DEGs) shared by at least four datasets. Values represent relative expression changes (log_2_FC) under warm temperature treatment (28°C vs 21°C). Top panel: core up-regulated DEGs. Bottom panel: core down-regulated DEGs. Common gene name for top marker genes is annotated on the right. Intersection with 5 or 4 datasets are indicated with purple or orange strip on the left side, respectively. Values represent log_2_FC and “ns” indicate non-significant (statistical threshold: |log2FC| ≥ 1 and padj < 0.05) in the corresponding dataset.

### Differential gene expression in seedling poorly represents organ-specific regulation

It is elusive how much DEGs identified in whole seedling represent transcriptional regulation in individual plant organs. Hence, we dissected temperature-treated seedlings to obtain root, hypocotyl, and cotyledon samples for gene expression analysis (Figure 3a). Our seedling and organ-specific datasets are directly comparable, given that the exact same experimental conditions were applied to both, i.e., 3 days at 21°C, then 4 days at 21°C or 28°C, under 16h light (70 μmol m^−2^ s^−1^ white light) and 8h in the dark cycles, using the same growth chamber (Percival CU-22L), and samples harvested 7 hours after the beginning of the light period (ZT7). Typical responses to warm temperature, such as root, hypocotyl, and petiole elongation, were confirmed under these exact experimental conditions before harvesting (Figure S1). We observed that gene expression regulation is largely dependent on the sample type, rather than temperature treatment, as evidenced by data distribution in principal component analysis (PCA) (Figure 3b). Root, hypocotyl, cotyledon, and whole seedling data showed distinct clusters, where root and cotyledon were the most distant, while seedling was relatively closer to cotyledon, evidencing that cotyledon biomass dominates whole-seedling samples. Significance threshold-free analysis (RRHO2) revealed broad concordance between seedling and organs (Figure 3c). However, although conventional DEG analysis (|log2FC| ≥ 1 and adjusted p-value < 0.05) corroborated the relative similarity between seedling and cotyledon, such DEG similarity appears limited to a few genes (Figure 3d and Supporting Table 1). We found that 1,119, 264, and 223 are up-regulated DEGs only in root, hypocotyl, and cotyledon, respectively, and similarly for 692, 226, and 181 down-regulated DEGs. Hence, 1,606 up and 1,099 down organ-specific DEGs were not identified in whole seedling RNA-seq. Furthermore, 67% (1,025) and 78% (399) of up-and down-regulated DEGs in whole seedling are not DEGs in root, hypocotyl, or cotyledon (Figure 3e). Altogether, our results show that analysis of gene expression regulation in whole seedling is confounded by differential regulation at organ level, possibly caused by compositional averaging and detection-threshold effects contributing to inefficient and incomplete overlap between whole-seedling and organ DEGs.

**Figure 3.**
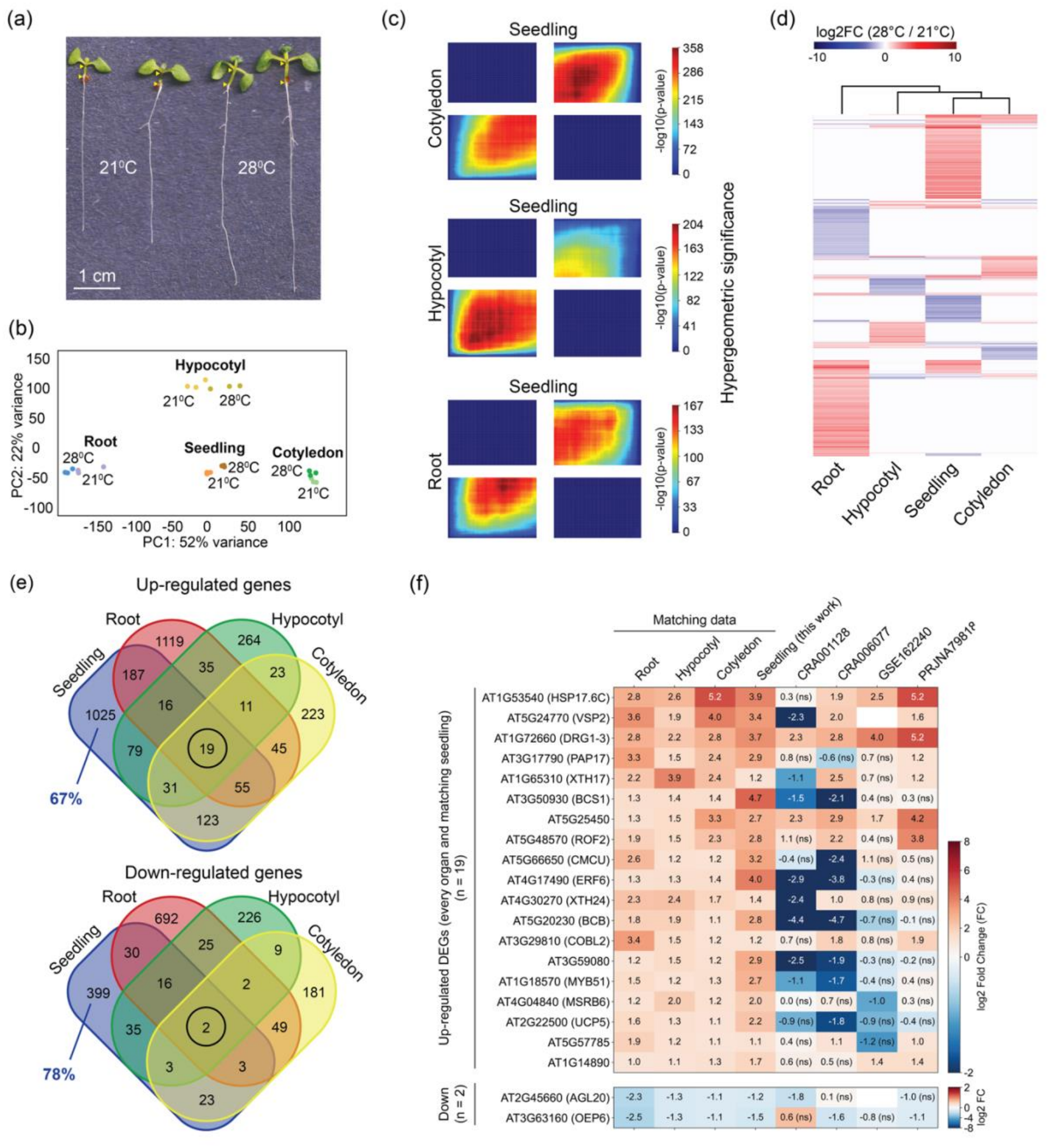
Organ-specific transcriptomic profile of thermomorphogenesis. **(a)** Representative phenotype of 7-day-old *A. thaliana* (Col-0) seedlings grown at continuous 21°C or transferred to 28°C for four days. Scale bar = 1 cm. **(b)** Principal component analysis (PCA) of transcriptomic profiles across whole seedlings and isolated organs (root, hypocotyl, and cotyledon) under 21°C and 28°C treatments. **(c)** RRHO2 heatmaps evaluating degree of gene expression concordance between whole seedlings and individual organs. Heatmaps depict threshold-free pattern agreement across entire ranked gene lists. Colour gradient indicates hypergeometric significance (-log_10_(p-value)), ranging from non-significant (blue) to highly significant (red). Quadrants represent concordant (upper-right: UP in both; lower-left: DOWN in both) and discordant (upper-left: UP in A, DOWN in B; lower-right: DOWN in A, UP in B) gene expression directionality. **(d)** Hierarchical clustering heatmap showing log_2_FC (28°C/21°C) of all DEGs identified across tissues and whole seedlings. **(e)** Venn diagrams displaying shared and unique up-regulated (top) and down-regulated (bottom) DEGs (|log_2_FC| ≥ 1.0, p_adj_ < 0.05) between whole seedlings and isolated organs. Percentages indicate seedling DEGs not detected as DEGs in any individual organ. **(f)** Heatmap of shared DEGs consistently regulated across all organs and whole seedlings. Color bar indicates log_2_FC (28°C/21°C).

**Supporting Figure 1.**
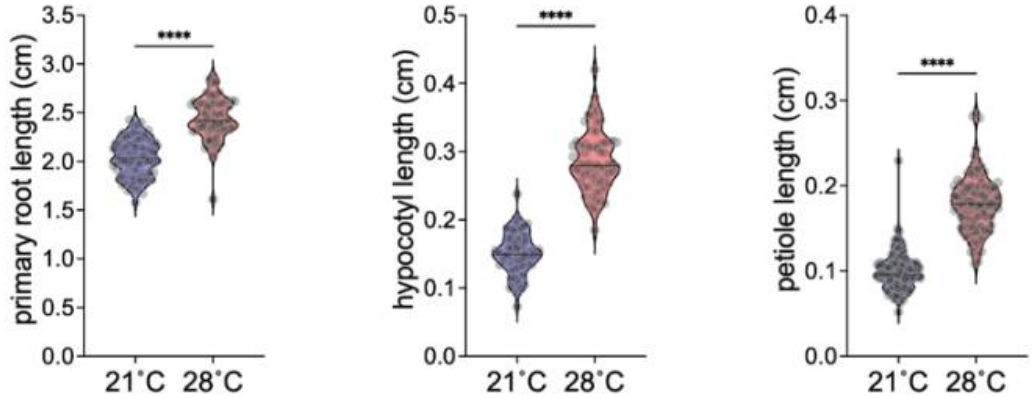
Quantified thermomorphogenic phenotype of 7-day-old *A. thaliana* (Col-0) seedlings grown at continuous 21°C or transferred to 28°C for four days. Temperature effect was calculated using two-tailed Student’s t-test (n > 65, ****p < 0.0001).

We identified 19 up and 2 down-regulated DEGs shared by every organ and matching whole seedling (Figure 3e-f). Among the up DEGs, we found *HEATSHOCK PROTEIN 17.6C* (*HSP17.6C*), suggesting a broader transcriptional activation for this *HSP* gene than others in the family. Similarly, broad up-regulation further included the transcription factors *ETHYLENE RESPONSE FACTOR 6* (*ERF6*) and *MYB DOMAIN PROTEIN 51* (*MYB51*), while *AGAMOUS-LIKE 20* (*AGL20*) was broadly down-regulated. We also identified *DRG1-3* and *AT5G25450* up-regulated in all organs, as well as in all seedling datasets (Figure 3f), suggesting a broad and robust induction by warm temperature treatments. Other genes were DEGs in most seedling datasets without evidence for discordant regulation (e.g., *ROF2*, *COBL2*, *AT1G14890*, and *AGL20*), while others showed discordant regulation in two or more seedling datasets (e.g., *MYB51*, *ERF6*, *BCS1*, *BCB*, and *AT3G59080*), consistent with the variability discussed above. Interestingly, other DEGs shared across all seedling datasets were differential only in specific organs, such as the heat stress-inducible *HSP70-HSP90 ORGANIZING PROTEIN 3* (*HOP3*) (up-regulated in seedling, hypocotyl, and cotyledon), and the cold stress-responsive *KIN1* (down-regulated in seedling, root, and hypocotyl) (Supporting Table 1). These results unveil previously unrecognised genes in the broad and organ-specific response to warm temperature regulated at transcription.

### Differential proteomics and regulated processes in response to warm temperature

Protein accumulation is often under the control of multiple regulatory layers involving mRNA transcription, translation, and stability, and protein stability. Hence, we produced organ-level differential proteomics in response to warm temperature treatment (Supporting Table 1 and 2). We identified 3810, 2725, and 4887 proteins in root, hypocotyl, and cotyledon, in which DEPs were 47 down/95 up, 158 down/27 up, and 53 down/104 up, respectively (Figure 4a). Hence, hypocotyl showed major suppression of protein accumulation, while root and cotyledon were biased towards increased accumulation, suggesting an organ-specific regulation of translation and/or protein degradation pathways. In fact, the bias in DEG direction was similar for root and cotyledon (i.e., roughly two-fold more up than down-regulated) but unrelated in hypocotyl, for which down/up ratio was 5.8 and 0.7 for DEP and DEG (Figure 4a-b), suggesting widespread transcription-independent regulation in hypocotyl. To ensure robust DEP and DEG comparison, we assessed the 20 genes with highest up or down DEP changes. These included various up-regulated proteins related to heat response, such as HSP90.4, DNAJ, HSP101, and CHAPERONIN-60ALPHA2 (Cpn60a2) across organs (Figure 4c). Furthermore, we observed apparent disconnect between differential protein and mRNA accumulation in every organ, with few cases indicating transcription as the main DEP driver. Among the top 20 down DEPs, we found reduction in several translation-related proteins in hypocotyl, including the EUKARYOTIC ELONGATION FACTOR 5A-2 (eIF-5A) and various ribosomal proteins (e.g., eL21z, uL1x, uL30w and uS4y). This is further supported by gene ontology (GO) analysis showing enrichment for ribosomal subunit among the top terms in down DEPs and, interestingly, without corresponding enrichment among DEGs (Figure S2). These results reveal a major suppression of protein accumulation in hypocotyl, likely driven by post-transcriptional inhibition of translational activity.

**Figure 4.**
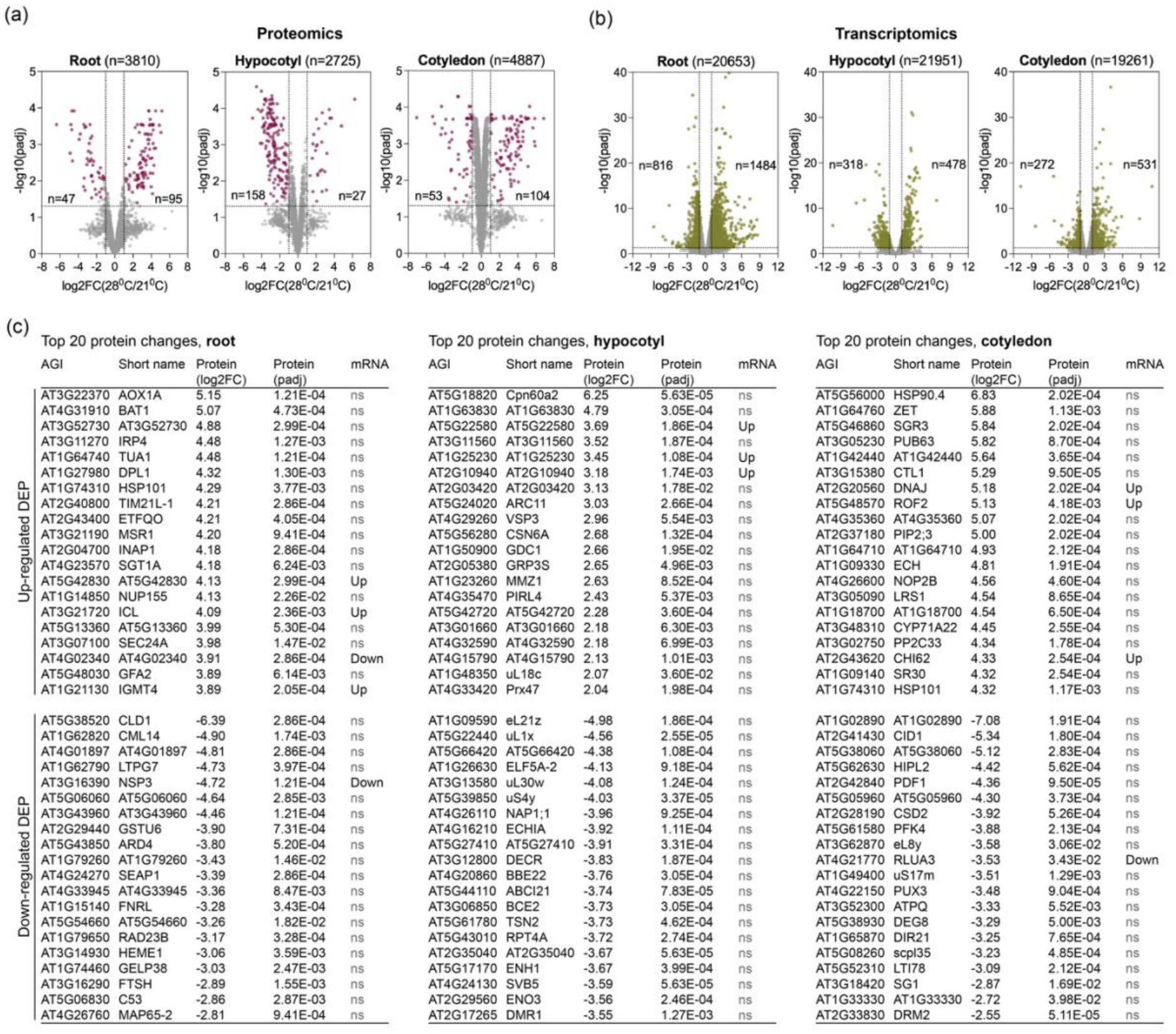
Organ-specific transcriptomic and proteomic changes in thermomorphogenesis. **(a-b)** Volcano plots displaying proteomic and transcriptomic differential abundance (28°C vs 21°C) in root, hypocotyl, and cotyledon. Dashed horizontal lines indicate significance threshold (p_adj_ < 0.05); vertical lines denote |log_2_FC| = 1.0 used selection of differentially expressed genes (DEGs). For differentially expressed proteins (DEPs), in addition to standard significance threshold (|log_2_FC| ≥ 1.0 and p_adj_ < 0.05), an imputation significance criterion was applied (see Method section). Total quantified features (n) and counts of significantly altered targets are annotated in each panel. **(c)** List of 20 genes with highest up or down DEP changes in root, hypocotyl, and cotyledon.

**Supporting Figure 2.**
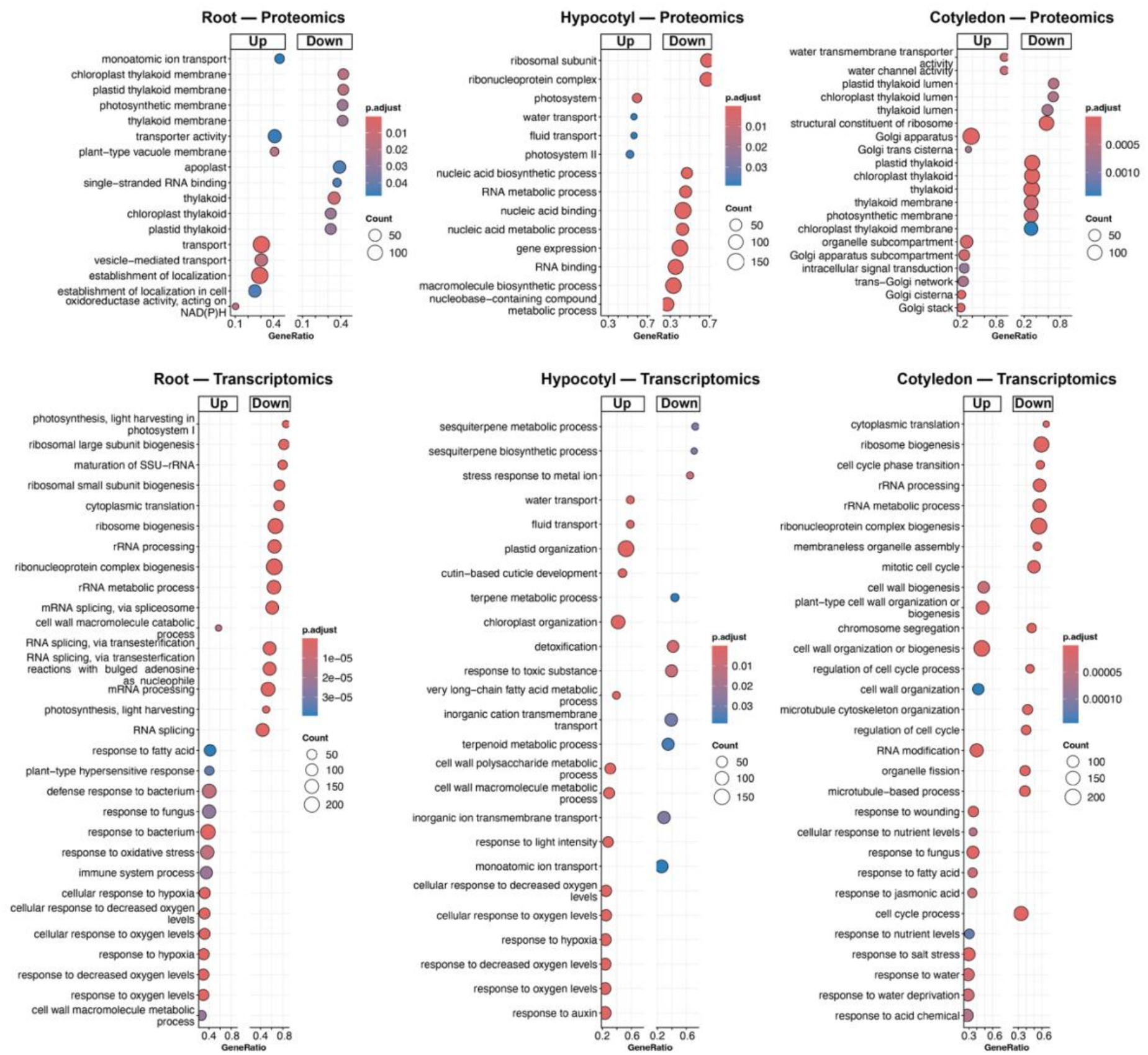
Gene ontology (GO) enrichment analysis comparing selected terms for up-(top) and down-regulated (bottom) biological pathways between transcriptomic and proteomics data across organs. Dot size corresponds to gene count, and dot colour indicates adjusted p-value.

Analysis of GO enrichment further revealed that proteomic changes were mainly associated with up-regulation in transport-related processes (e.g., water and vesicle-mediated transport) in root and cotyledon, as well as signalling in cotyledon, while changes in hypocotyl were mainly associated with down-regulation in RNA metabolism (Figure S2). Transcriptomic changes were associated with down-regulation in processes related to translation (e.g., ribosome biogenesis) and splicing in root; defence response (e.g. terpenoid metabolism) and ion transport in hypocotyl; and translation and cell division in cotyledon. Up-regulation was associated with processes related to response to pathogen and oxidative stress and cell wall metabolism in root; water transport, plastid, cell wall metabolism, and response to oxidative stress and auxin in hypocotyl; and cell wall metabolism, RNA modification, and response to pathogen and abiotic stress (e.g., nutrient levels, salt, and drought) in cotyledon. Therefore, our data show that translation is a central process under organ-specific thermomorphogenic repression at transcriptional level in root and cotyledon and post-transcriptional in hypocotyl.

### Widespread uncoupling of transcriptomic and proteomic changes in response to warm temperature

To further investigate the disconnect between DEPs and DEGs observed above, we produced a global comparison for genes found in both datasets. The correlation between transcriptomic and proteomic changes was poor in root and hypocotyl (Pearson correlation coefficient of 0.13 and 0.18, respectively) and only slightly higher in cotyledon (0.24) (Figure 5). We also found a limited number of gene overlap between DEPs and DEGs, i.e., 30, 13, and 12 in root, hypocotyl, and cotyledon, whereas 82%, 94%, and 93%, respectively, were DEPs without a statistically significant change in steady-state mRNA, suggesting major post-transcriptional regulation in every organ. Similarly, most DEGs were not accompanied by a statistically significant change in protein accumulation, suggesting transcriptional buffering. For instance, the major bias towards down DEP in hypocotyl was largely uncoupled from transcriptomic changes, further showing evidence for post-transcriptional regulation. Hence, our results support a global uncoupling between steady-state transcript and protein levels in response to warm temperature.

**Figure 5.**
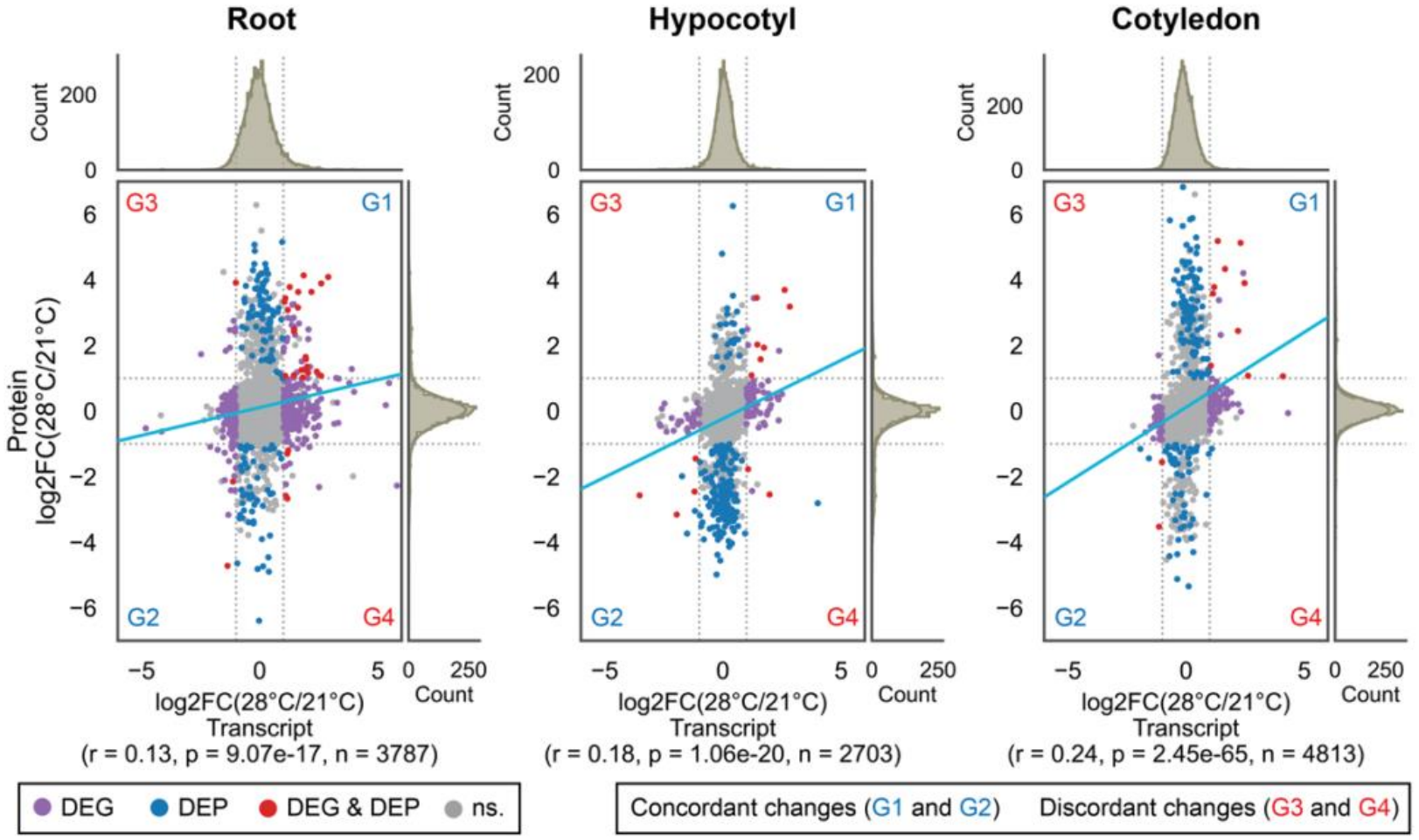
Comparison of transcriptomic and proteomic changes in root, hypocotyl, and cotyledon in thermomorphogenesis. Correlation analysis between transcript (log_2_FC) and protein (log_2_FC) responses per organ. Cyan lines represent linear regressions. Pearson correlation coefficients (r), p-values, and total overlapping features (n) are shown below each plot. Differentially expressed proteins (DEPs) and genes (DEGs) are in blue and purple, respectively, and overlapping DEP and DEG in red. Unchanged proteins or genes are in grey. Genes are group in quadrants (|log_2_FC| ≥ 1) for concordant (G1 and G2) and discordant (G3 and G4) changes.

### Shared transcriptomic and proteomic changes are mostly in the same direction

We next assessed the regulatory pattern in genes that are both DEPs and DEGs. We observed pronounced agreement in the regulatory direction (i.e., both DEP and DEG are up or down), i.e., 25 out of 30 genes in root, 11/13 in hypocotyl, and 12/12 in cotyledon (Table 1; see red dots in Figure 5). Among the concordant changes, 92% (23/25), 64% (7/11), and 83% (10/12) were up-regulated DEP and DEG in root, hypocotyl, and cotyledon, respectively. These genes are likely transcriptionally induced and post-transcriptionally stabilised or unaffected by warm temperatures. We found two concordant up-regulated genes shared by two organs, i.e., HSP90.1 (root and cotyledon) and AT5G25450 (root and hypocotyl). The latter was also found up-regulated in all whole seedling datasets and, although still uncharacterised, it might function as a ubiquinol–cytochrome-c reductase that has been shown to be upregulated in response to heat stress (Rizhsky *et al*., 2004). In root, we observed regulatory coordination to maintain elevated levels of, for instance, oxidative stress-related transcripts/proteins (GSTF6, GSTF7, TPX2, and DOX1), calcium homeostasis (ACA11), osmoregulation (OSM34), and tryptophan-derived secondary metabolism (IGMT1, IGMT4, CYP71B28, and CYP71B6), while concordant decrease was observed for the cold-regulated gene LTI78 and the nitrile-specifier NSP3. In hypocotyl, concordant increase included a low-affinity nitrate transporter (NRT1.11), a calcium binding protein (CCAP2), a putative acid phosphatase (AT1G25230), and a heat-induced gene (AT5G25450), while concordant decrease in a phosphate transporter (PHT1;1) and a cell wall modifying enzyme (EXPB3). In cotyledon, coordinated transcript/protein increase was associated, for instance, with heat response (HSP90.1, HSP70, DNAJ, and FKBP65), lipid metabolism (LACS3 and AT4G23680), and ER homeostasis (MEB1), while two RNA-binding proteins involved in cold response (GRP4) and RNA pseudouridylation (RLUA3) were concordantly reduced. Although we identified few discordant DEP and DEG changes, there is a case worth noticing. MSRB6 (METHIONINE SULFOXIDE REDUCTASE B6) is among the 19 core transcripts transcriptionally up-regulated across all organs and whole seedlings (Figure 3f), yet its protein level is significantly down-regulated in root and hypocotyl (Figure 5), suggesting a robust example of post-transcriptional regulation. Altogether, we identified several organ-specific genes displaying mostly concordant transcript and protein regulation, many of which uncharacterized in temperature response.

## DISCUSSION

Our work presents a comprehensive organ-specific transcriptomic and proteomic analysis of Arabidopsis seedlings grown under warm temperature. We showed that whole-seedling transcriptomics can present a distorted picture of temperature response, with 67% of up-regulated and 78% of down-regulated DEGs in seedling not showing differential regulation in root, hypocotyl, or cotyledon. We then presented organ-level transcriptomic and proteomic changes, unveiling regulatory processes and specific genes currently uncharacterized in the response to warm temperatures.

We observed that a third to half of reported DEGs in seedlings across comparable studies are study-specific, suggesting that DEG identification in thermomorphogenesis is sensitive to a combination of lab-to-lab variance and biological heterogeneity likely arising from differences in treatment duration, light regime, and circadian sampling. However, significance threshold-free analysis revealed broad concordance between the various whole seedling datasets, suggesting the existence of a core subset of genes in thermomorphogenesis. Organ-specific DEG analysis identified *DRG1-3* as up-regulated in every organ and whole seedling dataset. DRGs are a highly conserved family of GTP-binding proteins found in archaea, plants (three homologues in Arabidopsis), fungi, and animals (Bourne *et al*., 1991). Although plant DRGs are still poorly characterized, there is *in vitro* evidence that their GTP hydrolysis activity is activated upon heat stress, and a role as chaperone in ribosome assembly in response to temperature increases has been proposed (O’Connell *et al*., 2009). It is thus conceivable that translational regulation involves DRG1-3 in thermomorphogenesis.

Temperature increases promote fast hypocotyl cell elongation to produce taller embryonic stem—an energy-intensive process constrained by cell wall and regulated by water uptake and turgor-driven expansion (Cosgrove, 2024; Park *et al*., 2017). It is likely that the initial hormonal and transcriptional burst, cell wall loosening, and other major cellular changes to support expansion are followed by an elongation state that is sustained over days (four, in our experimental setup) through a global downscaling in energy-intensive processes, such as translation. Our work identified translational control as a likely central process in organ-specific thermomorphogenesis, particularly involving post-transcriptional/translational repression of translation machinery in hypocotyl. A previous work reported nearly equal number of genes with reduced or enhanced translation efficiency (693 down and 702 up) in whole seedlings (Chung *et al*., 2020), indicating that the effect we observed in hypocotyl is likely diluted in their data. Therefore, organ-specific studies on translational regulation have great potential to unveil key general repressive mechanisms involved in the sustained plant growth under warm temperatures. For instance, we identified MSRB6 among the most robust DEG upregulations (i.e., up-regulated across all organs and whole seedling datasets), yet its protein level is significantly down-regulated in root and hypocotyl—a clear case of mRNA and/or protein regulation, possibly involving translational control.

We identified the transcription factors *ERF6*, *MYB51*, and *AGL20* as DEGs in every organ and matching whole seedling, suggesting a broad regulation in response to warm temperature. However, *ERF6* and *MYB51* were up-regulated in our dataset but down-regulated DEGs in publicly available seedling datasets, further exemplifying the variability discussed above. *AGL20* (aka *SOC1*, *SUPPRESSOR OF OVEREXPRESSION OF CO 1*), on the other hand, was consistently down-regulated across seedling datasets, without evidence for discordant regulation. *AGL20*/*SOC1*, together with *FLOWERING LOCUS T* (*FT*) and *LEAFY* (*LFY*), are well-characterized targets of the master transcription factor CONSTANS (CO), in which CO induces their expression to promote flowering in long-day conditions (reviewed in Capovilla *et al*., 2015). *CO* is induced by GIGANTEA (GI), whereas CO protein is repressed by the photoreceptor phyB but stabilised by CRYPTOCHROME 2 (cry2) (Valverde *et al*., 2004). Warm temperature promotes flowering acceleration through PIF4 and PIF5 in a CO-independent manner in Arabidopsis (Thines *et al*., 2014). In *Osmanthus fragrans*, warm temperatures inhibit flowering through transcriptional repression of *AGL20* expression (Ye *et al*., 2025). We found *CO* down-regulated in two whole seedling datasets (this work and CRA001128, Jin *et al*., 2020) but unaffected in the organs, suggesting a weak contribution of CO towards *AGL20* expression. Our identification that *AGL20* is broadly down-regulated without clear association with flowering regulation at warm temperatures, suggests that AGL20 might antagonise thermomorphogenesis via other mechanism(s).

Within our dataset, we observed concordant DEP and DEG at organ-level for a limited number of genes. These genes are likely under sustained transcriptional regulation, rather than part of an initial temperature response. For instance, we observed concordant up-regulation related to oxidative stress and secondary metabolism in root. It has been shown that redox state regulation is required in various thermomorphogenic phenotypes, such as hypocotyl elongation (Dard *et al*., 2023; Raipuria *et al*., 2025) and growth in shoot apex and emerging leaves (Osborne *et al*., 2025), supporting a possible role for our identified oxidative stress-related genes in root thermomorphogenesis. Also in root, we found four key enzymes in tryptophan-derived secondary metabolism involved in glucosinolate biosynthesis with transcript and protein levels increased, i.e., IGMT1, IGMT4, CYP71B28, and CYP71B6. Tryptophan is a precursor for the auxin indole-3-acetic acid (IAA) and defence molecules indole glucosinolates (Barco and Clay, 2019). IGMT1 and IGMT4 are involved in the biosynthesis of methoxylated indole glucosinolates (Barco and Clay, 2019), while CYP71B6 metabolises indole-3-acetonitrile (IAN) to chemical defence compounds (Böttcher *et al*., 2014). IAN is at a metabolic crossroad between auxin and defence pathways, in which IAN can be hydrolysed into IAA by NITRILASE 1 (NIT1), NIT2, and NIT3 (Wójcikowska, 2026). Surprisingly, protein and transcript level of these nitrilases are not affected by temperature treatment in root in our data, suggesting that warm conditions channel tryptophan metabolism into the biosynthesis of indole glucosinolates and indolic defence compounds in root, potentially competing with auxin biosynthesis for tryptophan precursor.

In hypocotyl, concordant DEP and DEG included a cell wall modifying enzyme (EXPB3 downregulated), nutrient transporters (NRT1.11 upregulated and PHT1;1 downregulated), and a calcium binding protein (CCAP2 upregulated). Upon temperature increases, hypocotyl cells elongate to produce taller embryonic stem— constrained by the cell wall and regulated by water uptake and turgor-driven expansion (Cosgrove, 2024). Cell wall extension encompasses movement and separation of the cellulose microfibrils by a process of molecular creep, involving α-expansins (EXPA) induction of local dissociation and slippage of xyloglucans on the surface of the cellulose, and β-expansins (EXPB) possibly acting on different glycan for similar effect (Sampedro and Cosgrove, 2005). Our identification of EXPB3 downregulation suggests a selective usage of specific wall-loosening subfamilies, in which the β-expansin EXPB3 and perhaps others in the same family might act as antagonistic to hypocotyl elongation at warm temperatures. Moreover, rapid cell expansion during hypocotyl elongation at warm temperatures is an energy-intensive process, likely involving broad changes to nutrient assimilation, transport, and allocation. Our identification of NITRATE TRANSPORTER 1.11 (NRT1.11) upregulation and PHOSPHATE TRANSPORTER 1;1 (PHT1;1) downregulation, suggests changes in macronutrient transporter abundance that favour nutrient redistribution towards elongating hypocotyl, as previously shown for NRT1.11 in developing leaves (Hsu and Tsay, 2013). On the other hand, the identification of CYTOSOLIC CALCIUM-BINDING PROTEIN 2 (CCAP2) upregulation suggests a role for cytosolic calcium signalling in hypocotyl elongation under warm temperatures, as previously shown under other conditions (Folta *et al*., 2003; Li *et al*., 2011). Altogether, our findings on hypocotyl are broadly consistent with the literature and identified specific, possibly novel important players in thermomorphogenic stem elongation.

In cotyledon, concordant DEP and DEG included proteins involved in heat stress (upregulated HSP90.1, HSP70, DNAJ, and FKBP65), lipid metabolism (upregulated LACS3 and AT4G23680), ER homeostasis (upregulated MEB1), and RNA-binding proteins (downregulated GRP4 and RLUA3). In fact, cotyledon had the highest proportion of concordant upregulation of heat stress-related genes among the different organs. Furthermore, upregulation of LONG-CHAIN ACYL-COA SYNTHETASE 3 (LACS3) and AT4G23680, both involved in lipid metabolism, might be associated with changes in membrane fluidity and prevention of excess water loss, although AT4G23680 has also been previously identified among proteins localised to heat-induced stress granules (Kosmacz *et al*., 2019). MEMBRANE OF ER BODY 1 (MEB1), on the other hand, localises to endoplasmic reticulum (ER) bodies, i.e., rod-shaped organelles that accumulate β-glucosidases involved in plant defence against herbivores (Nakazaki *et al*., 2019) and facilitate root microbiota establishment in Arabidopsis (Basak *et al*., 2024). The function of MEB1 and its homolog MEB2 in glucosinolate metabolism is unclear but both proteins exhibit iron and manganese transport activity and are required for the rod-shaped morphology of ER bodies (Yamada *et al*., 2013). Given that warm temperatures promote growth over defence, it is possible that ER bodies are also affected, with MEB1 playing an uncharacterized role in leaf thermomorphogenic growth-defence tradeoff. We further identified two cases of concordant downregulation at transcript and protein steady-state levels: GLYCINE-RICH RNA-BINDING PROTEIN 4 (GRP4) and RLUA3, RNA-binding proteins (RBPs) involved in cold and salt stress response (Kwak *et al*., 2005) and enzymatic conversion of pseudouridine (Ψ) into uridine (U) (Li *et al*., 2025), respectively. Pseudouridylation is the most abundant modification type in tRNA, common in rRNA, and also found in plant mRNAs, for which a multilayered translational regulation has been described in Arabidopsis and rice (Li *et al*., 2025). Thus, the downregulation of RLUA3 and GRP4 shows that warm temperatures might post-transcriptionally regulate selected RNAs through changes to RNA modification and RNA-protein landscape, which has been shown in heat stress (Tong *et al*., 2022; Kan *et al*., 2023; Tang *et al*., 2020) but not in thermomorphogenesis.

The subset of genes showing concordant DEP and DEG in at least one organ and also found shared in seedling datasets was limited to eight cases: up-regulated *THIOREDOXIN-DEPENDENT PEROXIDASE 2*/PEROXIREDOXIN IIC (*TPX2*/*PRXIIC*), *HSP23.5*, *HSP90.1*, *DNAJ*, *HSP70*, *AT5G25450*, and *AT4G23680*, and down-regulated *LOW-TEMPERATURE-INDUCED 78*/*RESPONSIVE TO DESICCATION 29A* (*LTI78*/*RD29A*). These are genes likely under strict regulatory control in response to warm temperatures, consistent with the role of heat-shock proteins (HSP23.5, HSP90.1, DNAJ, and HSP70) and related (AT5G25450). *TPX2* is a concordant DEP and DEG up-regulated in root, as well as up-regulated in 4/5 whole seedling datasets, suggesting a prominent role for ROS sensing in the cytosol. Arabidopsis genome encodes for three peroxiredoxins (PRXIIB, C and D) that function as cytosolic thiol peroxidases ROS sensors (Vogelsang and Dietz, 2023). For example, PRXIIB has been shown to sense hydrogen peroxide (H_2_O_2_) upon immune activation, in which the *prxIIB* single and *prxIIb prxIIc prxIID* triple mutants were equally more susceptible to bacterial infection than the wild-type, showing that TPX2/PRXIIC and PRXIID has negligible contribution to stomatal immunity (Bi *et al*., 2022). It is thus possible that TPX2/PRXIIC is involved in ROS sensing under other conditions, possibly contributing to root thermomorphogenesis, given its up-regulation pattern observed here. AT4G23680 is a putative polyketide cyclase/dehydrase and lipid transport superfamily protein, previously identified among proteins localised to heat-induced stress granules (Kosmacz *et al*., 2019). Polyketides are a large and diverse group of secondary metabolites found in prokaryotes and eukaryotes, such as chalcones and stilbenes, involved in various processes, from plant immunity to crop quality (Flores-Sanchez and Verpoorte, 2009). The Arabidopsis genome has at least 33 genes annotated as polyketide cyclase/dehydrase and lipid transport superfamily proteins, all of which still uncharacterized. On the other hand, LTI78/RD29A has been extensively studied for its role in cold and dehydration response, possibly acting as a hydrolytic esterase (Kaur *et al*., 2025). Therefore, upregulation of AT4G23680 and downregulation of LTI78/RD29A represent potentially novel, robust gene regulations in thermomorphogenesis involving uncharacterized lipid (LTI78/RD29A) and polyketide (AT4G23680) metabolism.

Altogether, the comprehensive organ-specific resource presented in this work provides a basis for in-depth questions in plant response to warm temperatures that cannot be asked using whole seedling material.

## MATERIAL AND METHOD

### Plant growth conditions and treatments

*Arabidopsis thaliana* Col-0 wild-type plants were used in all experiments. Plants were grown on half-strength Murashige and Skoog (MS) medium (Duchefa, Haarlem, Netherlands; M0221) and 0.8% (w/v) phyto agar (Duchefa; P1003), without addition of sucrose, and cultivated at 21°C for three days (120 μmol m^−2^ s^−1^ white light), then at either 21°C or 28°C for another four days (70 μmol m^−2^ s^−1^ white light), under long day light cycles (16h light and 8h dark) throughout the entire experiment. Temperature treatment was performed using the same growth chamber (Percival CU-22L). Samples were harvested in triplicates at 7 hours after the beginning of the light period (ZT7).

### RNA sequencing analysis

RNA was extracted using Quick-RNA MiniPrep Kit (Zymo Research; R1054) and sent to Novogene (England) for mRNA isolation, library preparation, and sequencing on PE150. Sequencing reads obtained with our samples, as well as reads obtained from publicly available repository (Genome Sequence Archive, GSA, accession number CRA001128, CRA006077, GSE162240, PRJNA798184, and PRJNA1330348), were mapped and counted using the Nextflow pipeline nf-core/rnaseq (3.12.0) (Patel *et al*., 2025) against TAIR10.56 GTF, as reference annotation. Quantification of differentially expressed genes was performed using DESeq2 (Love *et al*., 2014).

### Proteomic analysis

Root and hypocotyl protein were isolated from material ground on liquid nitrogen using two volumes of lysis buffer (20 mM Tris HCl, pH 7.5, 0.5 M LiCl, 0.5% lithium dodecyl sulfate (LiDS), 0.4% IGEPAL CA630, 2.5% polyvinylpyrrolidone 40, 5 mM DTT, 10 mM ribonucleoside vanadyl complex, and 1.5X Roche EDTA-free protease inhibitor) per gram of powder after incubation for 30 min at 4°C, followed by debris removal via two sequential centrifugations at 10’000xg for 15 min. Protein in the cleared lysate was precipitated using four volumes of 100% acetone at -20°C for 1 h, followed by centrifugation at 20’000xg for 30 min at 4^°^C, then washed once with 100% acetone (incubation at -20°C for 1 h and pelleting by centrifugation). Cotyledon protein produced persistent aberrant peak drop in mass spectrometry using the extraction protocol described above, therefore, we used a previously described protocol optimised for leaf material (Buchs *et al*., 2018). For 50 mg of cotyledon powder, we added 0.2 mL of 50 mM Tris/HCl, pH 8.0, 4% (w/w) SDS, 50 mM DTT, and 1X Roche EDTA-free protease inhibitor. The mixture was vortexed and agitated for 30 min at 4°C. The suspension was centrifuged for 15 min at 16’000xg and 4°C. The supernatant was heated for 5 min at 95°C, cooled to room temperature before addition of one-fourth volume of 1 M iodoacetamide in 100 mM Tris/HCl, pH 8.0 and incubation for 30 min at 37°C. Protein was precipitated by addition of 10% (w/w) TCA and 0.07% DTT (w/w) in acetone by overnight incubation at −20°C, followed by centrifugation for 15 min at 16’000xg and 4°C. The supernatant was discarded and the pellet washed four times with ice-cold 0.07% (w/w) DTT in acetone. Protein pellets (root, hypocotyl, and cotyledon) were resuspended (8M urea in 100 mM Tris/HCl, pH 8), digested with trypsin, and peptides were isolated and analysed on nanoLC-MS/MS by the Core Facility Proteomics & Mass Spectrometry at the University of Bern, Switzerland. Data interpretation was performed using Spectronaut software operating in hybrid directDIA+ (Deep) mode. Protein inference was conducted based on the parsimony principle to determine the minimal list of proteins accounting for the detected peptides, controlling the false discovery rate (FDR) at 1%. Common contaminants (e.g., keratin, trypsin) were identified using a contaminant search database and removed prior to downstream differential abundance testing. For comparative and statistical analysis, fragment intensities were filtered (retaining proteins detected in the required minimum number of replicates per experimental group) and renormalised via median normalisation. Protein relative abundances were quantified using maxLFQ intensity values calculated with the R package *iq* (Pham *et al*., 2020). In parallel, protein abundances were assessed using the Top3 method, defined as the sum of the intensities of the three most abundant, consistently selected peptides per protein group across samples. Absolute quantification within samples was evaluated using Intensity-Based Absolute Quantification (iBAQ) values. Missing values were imputed to enable statistical comparisons. Protein imputation was performed using two distinct strategies depending on the metric. For iSN and iQN, proteins with at most one non-zero value within a group had their remaining missing values imputed using a left-censored Gaussian draw on a per-sample basis, drawing from a Gaussian distribution with a width of 0.3 times the sample standard deviation, centred at the sample distribution mean minus 2.5 times the sample standard deviation; any subsequent missing values were then imputed via Maximum Likelihood Estimation (MLE). For iTop3, peptide-level imputation was performed via Gaussian replacement on a per-sample basis, drawing from a distribution with a width of 0.3 times the sample standard deviation centered at the sample distribution mean minus 2.8 times the sample standard deviation. To ensure robustness, all imputation procedures were repeated 20 times, and the significance of each protein was recorded as a proportion of significance, being flagged if persistent across all cycles. Differential expression analysis to assess significant changes in protein abundance was performed using imputed Spectronaut intensity (iSN), while imputed IQ (iIQ) and Top3 (iTop3) are also provided in Supplemental Table 2 as an additional resource. We used PRIDE (Perez-Riverol *et al*., 2025) to deposit the mass spectrometry data to the ProteomeXchange Consortium.

### Rank-rank hypergeometric overlap (RRHO) analysis

Rank-Rank Hypergeometric Overlap (RRHO) analysis was performed to evaluate pattern agreement and overlap significance between differential expression datasets using the RRHO2 R package (Cahill *et al*., 2018). For each dataset, genes were ranked according to a metric combining statistical significance and expression directionality, calculated as -log_10_(p-value) x sign(effect size). RRHO2 objects were initialized with -log_10_ transformations enabled (log10.ind = TRUE) to compare gene lists across experimental conditions. Overlap hypermaps and stratified comparisons—including concordant (up/up and down/down) and discordant (up/down and down/up) expression patterns—were visualized via heatmaps.

### Gene ontology (GO) enrichment analysis

Gene Ontology (GO) functional enrichment analysis was performed using Gene Set Enrichment Analysis (GSEA) implemented via the clusterProfiler R package version 4.16.0 (Wu *et al*., 2021). Input gene lists were ranked by their mean log_2_(fold change) values in descending order. GSEA was conducted across all GO ontologies (Biological Process, Molecular Function, and Cellular Component) using *Arabidopsis thaliana* annotations (org.At.tair.db, TAIR key type) with gene set size bounds set to a minimum of 10 and a maximum of 500 genes. Statistical significance was adjusted using the Benjamini-Hochberg (BH) false discovery rate method with a threshold of p_adjust_ < 0.05. Enriched terms were categorized by regulation direction (activated/upregulated and suppressed/downregulated) and compared across tissue types (root, hypocotyl, and cotyledon) and experimental datasets (RNA-seq and proteomics) using ggplot2 (Wickham, 2016).

### Set intersection analysis

To evaluate shared and dataset-specific differentially expressed genes (DEGs) across multiple datasets, set intersection analyses were performed and visualized using UpSet plots in Python (pandas, matplotlib). Differentially expressed genes were categorized into up-regulated (log_2_FC ≥ 1.0, p_adj_ < 0.05) and down-regulated (log_2_FC ≤ -1.0, p_adj_ < 0.05) subsets for each dataset. Exact set intersections were calculated across all dataset combinations. Intersection sizes and combination matrix connections were plotted separately for up-and down-regulated genes, and high-order core gene sets present in at least four datasets (≥4) were extracted for downstream analysis. Venn diagrams were produced using the webtool Venn (https://bioinformatics.psb.ugent.be/webtools/Venn).

## DATA AVAILABILITY

RNA sequencing data generated in this work were deposited to the National Center for Biotechnology Information (NCBI) under the BioProject PRJNA1330348. The mass spectrometry proteomics data were deposited to the ProteomeXchange Consortium via the PRIDE partner repository with the dataset identifier PXD086939.

## AUTHOR CONTRIBUTIONS

RSR conceived and designed the project. JH produced all experimental data. SG performed RNA sequencing data analysis of our datasets. DJV performed RNA sequencing metanalysis. RSR performed additional bioinformatic analysis. RSR wrote the manuscript. All authors approved the manuscript.

## ACKNOWLEDGMENTS

We thank the Core Facility Proteomics & Mass Spectrometry at the University of Bern, Switzerland, for proteomic analysis.

## SUPPORTING INFORMATION

**Supporting Table 1.** Gene expression and simplified proteomics datasets.

**Supporting Table 2.** Proteomics dataset.

